# Fanconi Anaemia E3 Ligase complex activity is regulated by a druggable metabolite binding site in FANCX

**DOI:** 10.64898/2026.08.07.743451

**Authors:** Michael F Sharp, Yi Sing Gee, Jennii Luu, Karla J. Cowley, Henry Beetham, Christopher G. Langendorf, Jonathan S. Oakhill, John W Scott, Debora Cavero, Jordi Minguillon, Daqing Che, Jonathan B. Baell, Andrew J Deans, Jordi Surrallés, Kaylene J. Simpson, Wayne Crismani

**Affiliations:** St Vincent’s Institute of Medical Research, Fitzroy, Australia; The Faculty of Medicine, Dentistry and Health Science, The University of Melbourne; Medicinal Chemistry Theme, Monash Institute of Pharmaceutical Sciences, Monash University, Parkville, Australia; Victorian Centre for Functional Genomics, Peter MacCallum Cancer Centre, Melbourne, Australia; IR Sant Pau, Sant Pau Hospital, Barcelona, Spain; The Autonomous University of Barcelona (UAB), Spain; Advanced Therapies Mixed Unit, CIEMAT/IIS-FJD, Madrid, Spain; Zhejiang Jiuzhou Pharmaceuticals Inc.; 99 Waisha Road, Taizhou, Zhejiang, China; School of Pharmaceutical Sciences, Nanjing Tech University, Nanjing 211816, People’s Republic of China; Center for Biomedical Network Research on Rare Diseases (CIBERER), Barcelona, Spain; Sir Peter MacCallum Cancer Centre, Department of Oncology, University of Melbourne, Melbourne, Australia

**Keywords:** DNA repair, Fanconi anaemia pathway, drug discovery, AMPK, long chain fatty acyl-CoA

## Abstract

The Fanconi anaemia (FA) DNA repair pathway is an emerging target for precision cancer therapy. Using a high-throughput FANCD2-monoubiquitination assay, we identified a class of small molecules, including MSG010, that inhibit the FA E3 ligase complex *in vitro*. Because these molecules, and the metabolite, palmitoyl-CoA, are known to engage allosteric drug and metabolite (ADaM) binding site in AMP-activated kinase (AMPK), we hypothesised that a similar pocket exists within the FA complex. Supporting this, long-chain, but not short-chain, fatty acyl-CoA molecules inhibited the FA E3 ligase complex activity, and sequence analysis revealed similarity between the AMPK ADaM site and a WD40 repeat in the FA subunit FANCX. Targeted mutagenesis of this FANCX region disrupted E3 ligase activity or abolished inhibition by MSG010, suggesting the presence of an ADaM-like site in FANCX. Moreover, MSG010 preferentially killed *BRCA1*-deficient cells *in vitro*. These findings identify a putative small-molecule binding site in the FA pathway that may be developed further to test for exploitation as anticancer therapeutics.

## Introduction

The Fanconi anaemia (FA) DNA repair pathway is best known for its role in DNA interstrand crosslink (ICL) repair. Germline inactivation of the FA pathway leads to the condition called Fanconi anaemia, which results in predisposition to specific malignancies, progressive bone marrow failure, physical differences, infertility and other health issues^1–6^. Despite its role in rare disease, the FA pathway is also promising anti-cancer target given its central role in DNA interstrand crosslink repair and its intersection with homologous recombination^7–10^. A key step of the FA pathway is the monoubiquitination of FANCD2 and FANCI which is performed by the FA E3 ligase complex^11,12^. The monoubiquitination reactions clamp FANCD2-FANCI onto DNA and forms arrays around the site of DNA damage^13–18^. This process stabilises the DNA for downstream repair of the damage exclusively in S-phase and early G2 phase of the cell cycle^19,20^. Homologous recombination deficient cancer cells rely on the monoubiquitination of FANCD2 to stabilize DNA replication forks at sites of DNA damage^10,21^. Further, loss of the FA pathway has been suggested to be synthetic lethal with the loss of ATM^22^. This leads to an opportunity to target the FA E3 ligase complex to elicit a synthetic lethal response in multiple cancer types or perhaps function as a chemosensitising agent. Although numerous attempts have been made to identify small modulators of the FA pathway have been reported^23–29^, there is currently no FA E3 ligase complex inhibitors in the clinic.

The FA E3 ligase complex has only one known substrate, the FANCI-FANCD2 heterodimer. The E3 ligase complex consists of 9 proteins – FANCA, FANCB, FANCC, FANCE, FANCF, FANCG, FANCL and FANCX (formerly FAAP100) – along with the FA Associated Protein 20 (FAAP20)^12^. Central to the E3 ligase complex is FANCL, the E3 ligase, which is supported by FANCB and FANCX in forming a dimeric heterotrimer^11^. FANCX has been shown to be essential for E3 ligase activity, however, the functional role of FANCX is not known other than it structurally supports FANCL alongside FANCB^12,30,31^.

Using a reconstituted active FA E3 ligase complex with recombinant proteins^32^ we can assay for its activity via the detection of monoubiquitinated FANCD2 in a high-throughput assay^8^. We have used this assay to screen a library of 45,784 compounds to identify inhibitors of the FA E3 ligase complex. Through multiple rounds of secondary screening, we have found that a class of small molecules and long chain fatty acid CoA metabolites that inhibit the FA E3 ligase activity via allosteric inhibition of FANCX *in vitro*. The small molecules and long chain fatty acyl CoA metabolites are known activators of AMPK complexes containing the (β1-isoform) and interacts with a well-characterised Allosteric Drug and Metabolite (ADaM) binding site in AMPK that shares sequence homology with a region on FANCX^33^.

## Results

We used a high-throughput screening approach to identify inhibitors of the FA E3 ligase complex using the recombinant FANCD2 monoubiquitination assay described previously^8^. We screened over 45,000 small molecules in singlet reactions (Sup Fig 1). We set a z-score cut-off of −2 to select 1,332 compounds for secondary screening. Secondary screening was performed by using the recombinant FANCD2-monoubiquitnation assay with quadruplicate data points. From secondary screening, 305 compounds had an average z-score of less than −2, which were selected for a dose response assay. The dose-responses were performed with the recombinant FANCD2-monoubiquitnation assay over a 5-point dose range (0.2 - 20 µM). To rank compounds on potency, area under the curve (AUC) was calculated from the dose response curves (Sup Fig 1). From the screen, a known small-molecule activator of AMPK, A-769662, was identified as a preliminary hit and we chose to focus further on this molecule (primary screen z-score = −2.13, secondary screen z-score = −2.11, dose response AUC = 583). A-769662 has previously been shown to bind an <u>a</u>llosteric <u>d</u>rug <u>a</u>nd <u>m</u>etabolite binding site in the AMPK β1 complexes (ADaM site)^33^. The AMPK activator SC4 and many of its analogues also bind the ADaM site to activate AMPK^34,35^. We therefore determined if other AMPK activators could inhibit FANCD2 monoubiquitination. A panel of eleven SC4 analogues along with five other known AMPK activating compounds were analysed with the FANCD2-ubiquitination assay, over a 1.25 – 40 µM dose-response range. MSG010, a SC4 analogue, (Fig 1A) was the most potent of these compounds with an IC50 in the ubiquitination reaction of 8.4 µM. All but four of the AMPK activating compounds reached 50% inhibition in the dosage range of 1.25 – 40 µM. The FANCD2-ubiquitination cascade was then visualised via a western blot of the recombinant FANCD2 ubiquitination assay where the blot was probed for biotinylated ubiquitin. A slight reduction of FANCD2 ubiquitination was seen at 56 µM and 100 µM of MSG010 (Fig 1B). However, autoubiquitination of UBE2T was not inhibited by MSG010, unlike for the UBE1 inhibitor - TAK243 (Fig 1B) – suggesting that MSG010 inhibits the ubiquitination cascade via E3 ligase activity.

**Figure 1.**
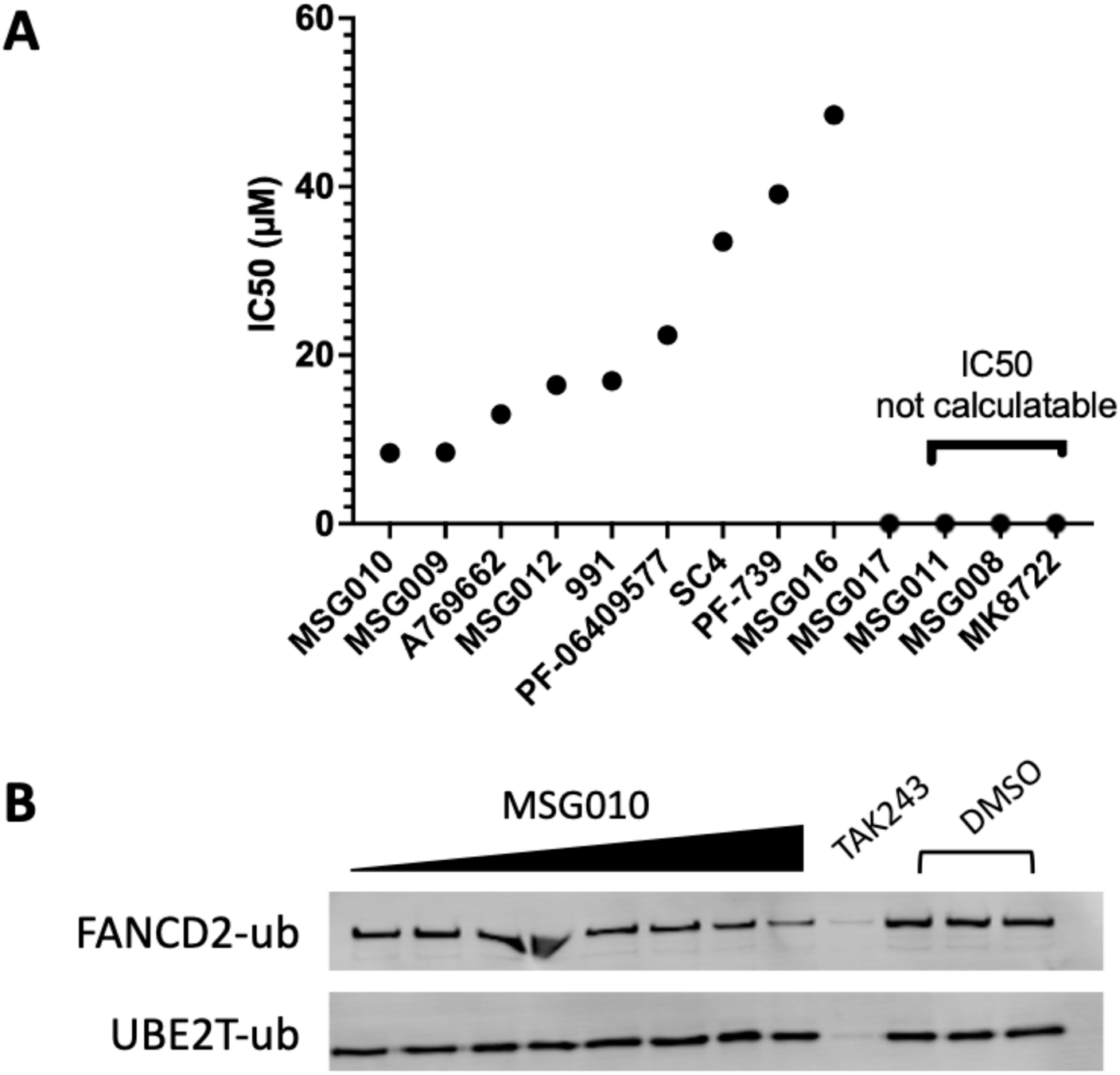
A) IC50’s calculated from the dose responses of 13 AMPK activating compounds in the recombinant FANCD2-monoubiquitination assay. B) Western blot of the recombinant FANCD2-monoubiquitination assay with a MSG010 dose response. The blot is probed for biotinylated-ubiquitin. The panels show mono-ubiquitinated FANCD2 and auto-ubiquitinated UBE2T. Controls were DMSO and the UBE1 inhibitor, TAK243.

Endogenous metabolites that activate AMPK via the ADaM site have been reported to included long chain fatty acyl-CoA’s ^33^. A set of fatty acyl-CoA’s were analysed with the recombinant FANCD2-ubiquitination assay to determine if they elicit a similar inhibitory effect to the small molecule AMPK activators (Table 2).

Fatty acyl-CoA’s with saturated acyl chain lengths ranging from 8 to 18 carbons were used in the FANCD2 ubiquitination assay (Fig 2A). The most potent inhibitors were the long chain fatty acyl-CoA’s myristoyl- (14:0), palmitoyl- (16:0) and stearoyl-CoA (18:0) (Fig 2B). Lauryl-CoA (12:0) exhibited markedly lower efficacy, while octanoyl-CoA (8:0) failed to inhibit the FA E3 ligase. Collectively, these data highlight an important structure-function relationship between acyl chain length and FA E3 ligase activity. Neither free Coenzyme A nor palmitic acid inhibited the reaction. Additionally, the saturation status of the fatty-acyl chain did not affect inhibition potency as palmiteoyl-CoA (16:1) elicited a similar inhibition profile to palmitoyl-CoA (16:0). The thiol-ester bond between the acyl chain and coenzyme A can be hydrolysed and result in non-enzymatic acylation of proteins *in vitro*^36^. To demonstrate that the inhibition observed was due to non-covalent binding and not covalent acylation, we tested a non-hydrolysable analogue, palmitoyl-ether-CoA. This showed similar results to inhibition of the reaction by palmitoyl-CoA.

**Figure 2.**
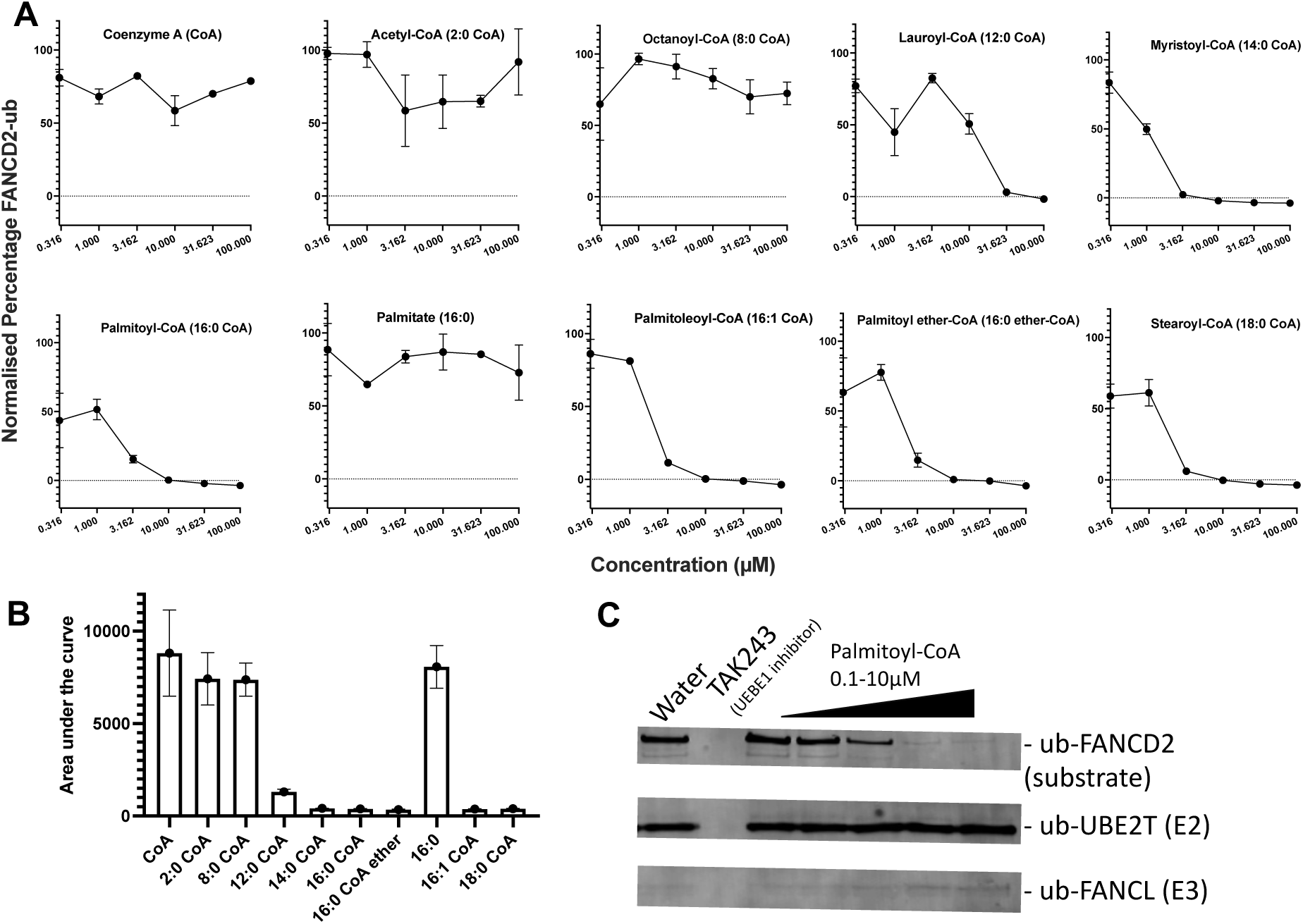
Analysis of fatty acyl-CoA in the recombinant FANCD2 ubiquitination assay. A) Dose responses of fatty-acyl-CoA’s in ubiquitination assay, which include 0, 2, 8, 12, 14 16 and 18 carbon chains, along with free fatty acid palmitate, unsaturated fatty acyl-CoA, and unhydrolysable fatty acyl-ether-CoA. B) Area under the curve calculated from the dose responses. The smaller the value, the more potent the inhibitor is. C) Western blot of the recombinant FANCD2-monoubiquitination assay with a MSG010 dose response. The blot is probed for biotinylated-ubiquitin. The panels show mono-ubiquitinated FANCD2, auto-ubiquitinated UBE2T and auto ubiquitinated FANCL Controls were DMSO and the UBE1 inhibitor, TAK243.

The inhibitory effect of palmitoyl-CoA on the ubiquitination of FANCD2 cascade was visualised by probing for biotinylated ubiquitin on a western blot of the recombinant FANCD2 ubiquitination reaction. Palmitoyl-CoA inhibited FANCD2 ubiquitination in a dose dependent manner but did not reduce auto-ubiquitination of UBE2T or FANCL (Fig 2C). This is consistent with the inhibition occurring at the E3 ligase step of the ubiquitination cascade, similar to the AMPK activator, MSG010 (Fig. 1B).

The SC4 analogues and palmitoyl-CoA inhibition of the FA core complex suggest the existence of a binding pocket for fatty acyl-CoA metabolites within the FA complex, similar to the ADaM site in AMPK. To identify which proteins within the FA E3 ligase complex these ligands interact with, the amino acid sequence of the surface of the human AMPK β1 ADaM site (V81 to F112) was aligned with the sequences of FA core complex proteins. Notably, the AMPK ADaM site shares high sequence identity to a region of the WD40 domain of FANCX with 8/32 exact matches and a further 6/32 amino acids with similar properties (Fig 3A). To identify important residues in this motif on FANCX, the alternative AMPK β isoform sequence (β2) was aligned with AMPK β1 and FANCX. SC4 analogues are less effective at activating AMPK β2 complexes than β1, and palmitoyl-CoA has no effect on AMPK β2 complexes^33^. This suggests that the residue similarities between AMPK β1 and FANCX, which are different to AMPK β2, may have an important role in FANCX interacting with these molecules.

**Figure 3.**
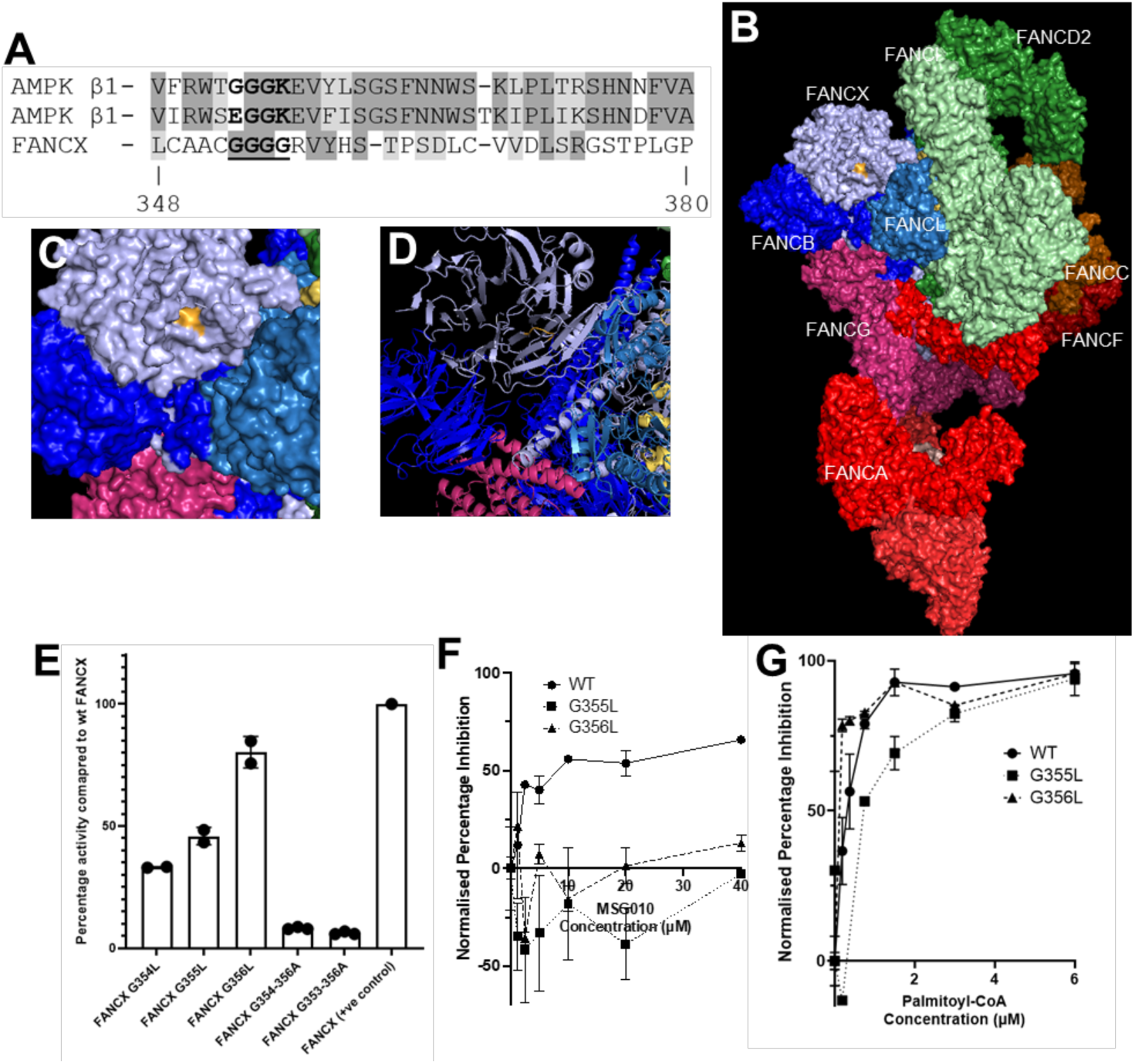
Proposed compound binding site on FANCX. A) Alignment of the ADaM site in the AMPK β1 and β2 subunits and FANCX. B) Structure of the FA core complex bound to the FANCD2-FANCI heterodimer (image generated in PyMol based on PDB 7KZD) C) The four glycines in the ADaM site are situated in the beta turn of one of the WD40 repeats in the WD40 domain of FANCX. D) In gold, the four glycine residues predicted to be involved in the ADaM site, which are accessible on the surface of FANCX and are spatially situated near the active ubiquitin ligase, FANCL. E) Activity of the FA core complex in the recombinant FANCD2 monoubiquitination reaction, with the variants of the glycines situated in the ADaM site. F) Dose responses of MSG010 in the recombinant FANCD2 monoubiquitination reaction in presence of G355L and G356L mutations demonstrating the inhibitory effect is lost with the mutant FANCX. G) Dose responses of Palmitoyl-CoA in the recombinant FANCD2 monoubiquitination reaction in presence of G355L and G356L mutations demonstrating the inhibitory effect is not lost with the mutant FANCX’s.

Only 1/8 of the identical residues differed between FANCX-AMPK B1 and AMPK β2 which was FANCX G353. G353 is the first residue of a glycine tetrad (G353-356) in which the first three are shared with AMPK β1. The four glycines are positioned in one of the beta-turns of the FANCX WD40 domain. The glycine tetrad on FANCX (Fig 3A) is located on the surface of the protein (Fig 3B, C, D) making it accessible for the ligands. The four glycines were individually mutated to leucines in FANCX and expressed as recombinant proteins. The G353L mutant could not be expressed. The activity of the three remaining FANCX variants were tested in the recombinant protein FANCD2 monoubiquitination assay. All of them had reduced activity (Fig 3E) with G354L, G355L and G356L having 33%, 46% and 80% of the wild-type activity, respectively. Three more mutants were attempted to be expressed; G353-356A, G354-356A and G353-356del. The G353-356del could not be expressed and the G353-356A and G354-356A mutants had less than 7% activity in the FANCD2 monoubiquitination assay.

We hypothesised that these glycines in FANCX may have a role in the binding of the SC4 analogues and long chain fatty acyl-CoA’s. We tested this hypothesis by dosing the FANCD2-ubiquitination assay with MSG010 and palmitoyl-CoA in the presence of the two most active FANCX mutants, G355L and G356L (Fig 3F-G). The ability of MSG010 to inhibit the reaction was reduced in the presence of both FANCX mutants suggesting that G355 and G356 are important for ligand binding. Inhibition of the FA core complex by palmitoyl-CoA was not hindered by the G355L and G356L mutations. However, it is possible that a single point mutation of residue lacking a functional R-group may not change the binding of a long flexible molecule such as a long chain fatty acyl-CoA. Testing this further was precluded by the low activity of the expressible multiple-mutation FANCX variants in the recombinant FANCD2 ubiquitination assay.

Some of the upstream FA pathway genes have been reported to have a synthetic lethal relationship with *BRCA1* deficiency^10,21,37,38^. Therefore, we proceeded to test if MSG010 could preferentially kill *BRCA1-*deficient cells. MSG010 demonstrated a synthetic lethal effect in *BRCA1* isogenic RPE *p53-/-* cells at the highest doses (1 - 10 µM) where it reduced the growth of the *BRCA1*^-/-^ cells compared the *BRCA1^+/+^* (Fig 4A). The PARPi inhibitor olaparib was used as control (Fig 2B) for synthetic lethal killing of *BRCA1*-defcient RPE cells^39^.

**Figure 4.**
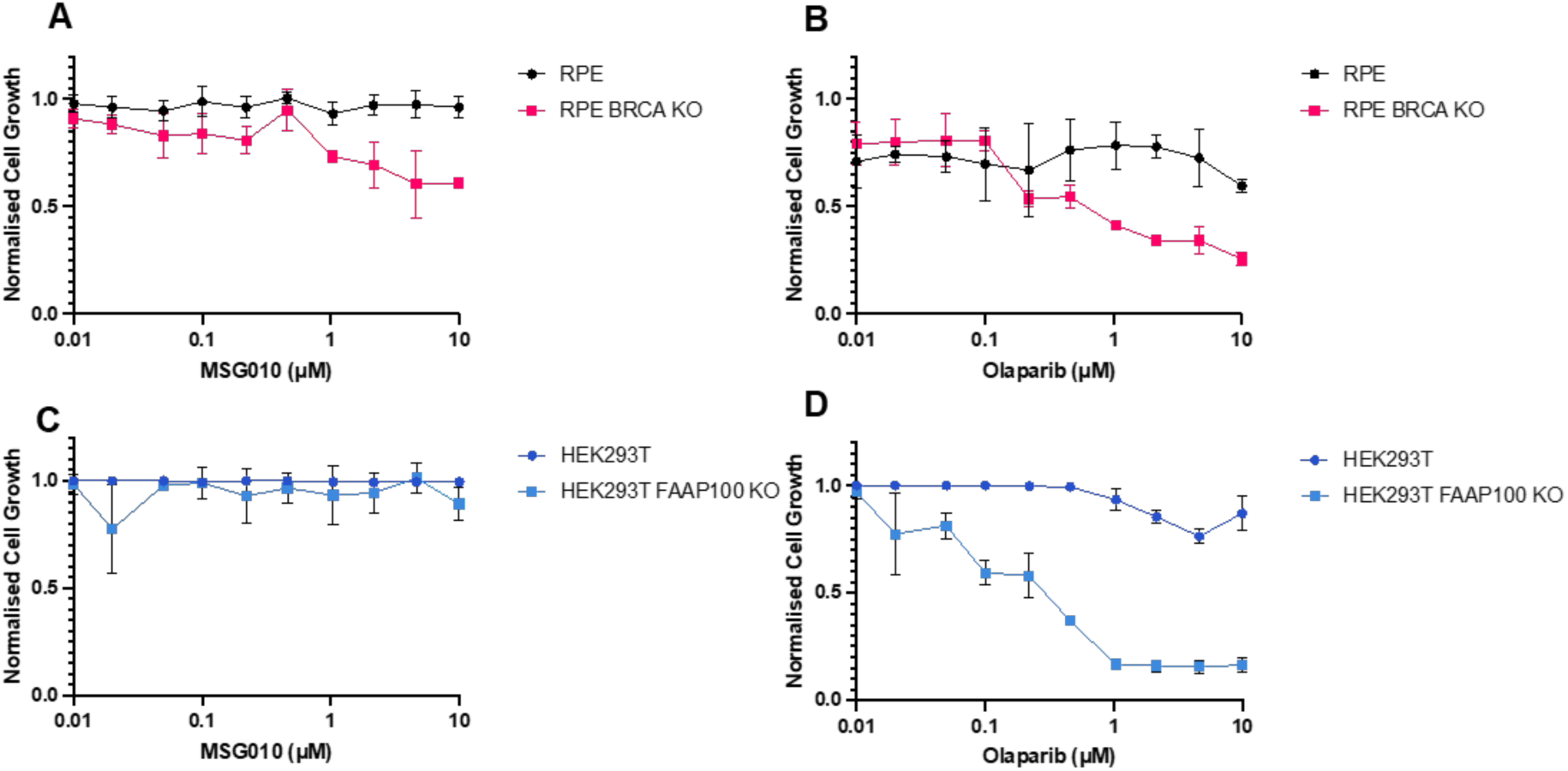
Cell growth assay with MSG010 dose response to assess synthetic lethal and pathway specificity. Cells were drugged at 24 h and 72 h, cells were counted at 144 h. Dose response was performed in two isogenic cell lines, RPE *p53*-/- with and without *BRCA1* to assess synthetic lethality (A and B) and HEK293T with and without *FANCX* to assess if the compound is targeting the FA pathway (C and D). Error bars represent standard error of the mean of three technical replicates. Error bars are S.E.M. mean of three independent experiments..

We generated *FANCX* knocked out HEK293T cells to attempt to identify target specific effects of MSG010 in the absence of its proposed target, however we found that unlike in RPE cells, both WT and *FANCX* knockout HEK293 were fully resistant to MSG010 even at the highest dose tested (Fig 2C). The *FANCX*-deficient cells were also sensitive to PARP inhibition (Fig 2D). MSG010 selectively inhibited the growth of BRCA1-deficient RPE cells. However, because both WT and *FANCX*-deficient HEK293T cells were resistant to MSG010, we were unable to determine whether this effect depended on *FANCX*. Further, we were unable to demonstrate that MSG010 causes FA-specific cellular phenotypes, as MSG010 did not reduce FANCD2 foci in HCT116 cells – a biomarker of FA E3 ligase activity (Sup Fig 2). HCT116 cells were used for the FANCD2 experiments because they have a high baseline level of FANCD2 foci^27^.

Finally, micronucleus and G2 arrest assays were performed as complementary approaches that can measure inactivation of the FA pathway^28^. In these two assays, loss of the FA pathway results in an increase of micronucleus formation and an increase of cells arresting in the G2 phase of the cell cycle when treated with diepoxybutane. A-769662 treatment did not phenocopy loss of the FA pathway using the micronucleus and G2 arrest assays (Sup Fig 3). In summary, recombinant protein assays identified A-769662 and MSG010 as candidate FANCX inhibitors, however neither compound produced convincing evidence of FA pathway inhibition in cell-based assays.

## Discussion

The discovery and development of DNA repair inhibitors for use in precision oncology is vital for the progression of cancer therapy. Various efforts have been made to identify FA core complex inhibitors due to the possibility of overcoming acquired tumour chemoresistance and the suggestions that loss of function of the FA core complex may be synthetic lethal with DNA repair and checkpoint deficiencies^10,21,22,40–42^. We screened over 45,000 compounds with a high-throughput recombinant protein FANCD2 monoubiquitination assay and identified A-769662, a palmitoyl-CoA mimetic drug and known AMPK activator, as a novel inhibitor of the FA core complex *in vitro*.

This led us to pursue the possibility that long chain fatty acyl CoA’s can also inhibit the FA core complex *in vitro*, possibly through a common allosteric drug and metabolite binding site. The MSG compounds were first developed as AMPK activators capable of selectively activating AMPK with distinct beta subunit isoforms, unlike AMPK pan-activators (e.g. MK8722)^35^. Among these, MSG010 and MSG011 are the most potent AMPK beta1 activators^35^. Interestingly, MSG010 was the most potent FA core complex inhibitor in this set of compounds. Further, MSG011 had a modest effect, but did not inhibit the FANCD2 ubiquitination assay by 50% at the maximum concentration used in our assay, 50 µM. Subsequently, MSG011 was one of the weaker FA core complex inhibitors in this set of compounds. The difference between the activity of MSG010 and MSG011 in our study provides the starting point to perform structure-activity-relationship studies to generate a compound that is selective for the FA core complex over AMPK.

MSG010 elicited a mild preferential killing of *BRCA1*-deficient RPE *p53*^-/-^ cells compared to isogenic *BRCA1*-proficient RPE *p53*^-/-^ at concentrations between 1 and 10 µM. Further, MSG010 had no effect on *FANCX* KO HEK293T’s which is the proposed target of the compound within the FA core complex. The inability for MSG010 to have an effect on the *FANCX* KO cells is consistent with the hypothesis that the compound does not cause non-specific DNA damage resulting in interstrand cross linking or replication fork stalling, as a *FANCX* KO cell would be expected to be sensitive to these types of damage^10,43–45^. However, the inability for MSG010 to reduce FANCD2 foci and induce micronuclei means we cannot conclude that the synthetic lethal effect is completely due to MSG010 inhibiting the FA core complex. This could be due to the compound having other known targets in the cell, namely, AMPK. AMPK is a key regulator of cell metabolism, switching anabolic processes for catabolism^46^. Activation of AMPK could result in numerous possible outcomes for the cell cycle and DNA repair which could affect *BRCA1* KO cells. It has been demonstrated that AMPK activates 53BP1 by phosphorylating Ser1317, resulting in the recruitment of canonical DNA non-homologous end joining (c-NHEJ) machinery to repair double strand breaks^47,48^. The use of c-NHEJ in S-phase to repair double strand breaks is error prone and can be catastrophic in cells with defective homologous recombination repair, such as in RPE *p53*^-/-^*BRCA1*^-/-^ cells^49^. Further investigation into the nature and extent of the inhibition of the FA core complex by MSG010, compared to its AMPK activation ability, needs to be explored before the preferential killing of *BRCA1-*deficient cells observed in this study can be attributed to FA core complex inhibition.

Our data support the hypothesis that long chain fatty acyl-CoA’s might inhibit the FA core complex *in vitro* in a drug like manner (not via acylation). Long chain fatty acyl-CoA’s are not only intermediates of both fatty acid synthesis and beta-oxidation but also function as signalling molecules that allosterically activate AMPK, signalling for the cells to produce more ATP ^33^. Currently, the physiological significance of the interaction between the FA core complex and long chain fatty acyl-CoA’s is poorly understood. The FA pathway is best known for its role in the repair of DNA interstrand cross-links, from both endogenous and exogenous sources^50^. The most likely endogenous source of the interstrand crosslinks which leads to bone marrow failure in individual with FA are reactive aldehydes^51–55^. The FA pathway is seen as the second tier of protection against DNA damage caused by aldehydes, the first tier being aldehyde dehydrogenase (*ALDH2*)^56,57^. A potential interaction of long chain fatty acyl-CoA’s and the FA core complex may be related to the DNA damage caused by aldehydes derived from metabolism. It could be possible that long chain fatty acyl-CoA’s inhibit the FA pathway in cells when the energy levels are not favourable for the cell cycle, functioning as a metabolic checkpoint to prevent the pathway from activating outside of S/G2 phase of the cell cycle. This is in line with the function of AMPK and its ability to be activated by long chain fatty acyl-CoA’s^33,46^. Without further experimentation which is beyond the scope of this study, we cannot conclude the inhibition seen in our recombinant FANCD2-ubiquitination assay represents what occurs in cells. However, this study has shown that long chain fatty acyl-CoA’s can have a drug-like effect on the FA core complex proteins *in vitro*.

FANCX is an integral protein in the FA core complex for FANCD2 monoubiquitination, but its exact function is unknown^58,59^. Two studies have shown that germline biallelic loss of *FANCX* results in Fanconi anaemia and that pathogenic variants lead to reduced FANCD2-monoubiquination^60,61^. The protein exists in a subcomplex with FANCB and the E3 ligase, FANCL, in a dimeric heterotrimer ^31^. FANCB has a role as a structural protein in which its dimer is the central scaffold of the FA core complex. Previous studies suggest FANCX could play a similar – yet non-redundant – role to FANCB due to its structural similarities, despite having little sequence homology with FANCB^30,31^. The proposed ADaM site on FANCX suggests that FANCX may have a greater function than as just a structural protein. The FANCX ADaM site is positioned on the WD40 domain. We investigated a glycine tetrad due to their sequence identity with the AMPK β1 ADaM site^33^. The importance of these glycines in AMPK β 1 (G86-88) was demonstrated by mutating G86E which mimics the AMPK β2 sequence. Mutation of this single glycine in AMPK beta1 is sufficient to render the kinase insensitive to allosteric activation by either A-769662 or palmitoyl-CoA. AMPK β1 G86 is the same residue that has sequence homology with the FANCX ADaM site but differs from AMPK β2. Due to the lack of side chain on the glycines it is unlikely that they directly interact with the ligands. The exact residues that mediate ligand binding are yet to be elucidated. One possibility is that the glycines confer flexibility near the binding site, facilitating ligand binding, or more generally contribute to correct protein folding.

The lack of response to the AMPK activators on the FA pathway in cell culture – despite supporting biochemical *in vitro* evidence – could be due to the compound being metabolised in the cell or the FA pathway effect being dampened by the compound’s activation of AMPK. Further, the metabolic regulation of the FA pathway may not be observable in high-energy *in vitro* cell culture conditions. However, *in vivo* there may be more nuanced conditions that give biological meaning to this interaction.

## Conclusion and future perspectives

The ability to perform SAR studies on the compounds described in this manuscript and compare this to previously published data on their interaction with AMPK β1 is promising for the development of a first-in-class specific FA core complex inhibitor. Further, the detailing of an allosteric drug and metabolite binding site on the FA core complex protein, FANCX, is intriguing and will lead to further investigation into interactions of the FA pathway outside of its canonical DNA repair role.

## Acknowledgements

Thank you to Susanne Ramm for helpful discussions. This work has been supported by funding from the National Health and Medical Research Council (Career development fellowship GNT1129757 to W.C., a project grant GNT1156343 awarded to W.C., A.J.D. and K.J.S.), Maddie Riewoldt’s Vision (MRV Fellowship WC-MRV2016 to W.C. and Grant-In-Aid SVI-MRV2017G to W.C. and A.D.), the Victorian Cancer (Victorian Cancer Agency Fellowships to W.C. and A.J.D.) NHMRC (GNT1129757, GNT1156343), the Victorian government IOS program, the Therapeutics Innovation Australia (TIA) Pipeline accelerator (awarded to M.F.S) and the St Vincent’s Institute of Medical Research Foundation Rising Star Fellowship and Award (Awarded to M.F.S). The Victorian Centre for Functional Genomics (VCFG) (K.J.S.) is funded by the Australian Cancer Research Foundation (ACRF), Phenomics Australia through funding from the Australian Government’s National Collaborative Research Infrastructure Strategy (NCRIS) program, the Peter MacCallum Cancer Centre Foundation and the University of Melbourne Research Collaborative Infrastructure Program. We thank Compounds Australia at Griffith University for their provision of specialized compound management and logistics services to the project. Griffith is a recipient of Queensland Government “Smart State Research Facilities Fund” funding and Australian Government funding provided under the “Super Science Initiative” and financed from the Education Investment Fund. The authors acknowledge the facilities, and the scientific and technical assistance of the Victorian Centre for Functional Genomics at Peter MacCallum Cancer Centre. JS team is funded by Ministerio de Ciencia e Innovación (PID2021-122411OB-I00/AEI/10.13039/501100011033/FEDER, UE; PID2024-155624OB-I00; PDC2022-133233-I00; PDC2025-165867-I00); Instituto de Salud Carlos III (ISCIII) and fondos Next Generation EU (ICI22/00076); ICREA-Academia program, Fundació Institut Català de Recerca i Estudis Avançats, Agència de Gestió d’Ajuts Universitaris i de Recerca (2021-SGR-00835, 2023-2025); Centro de Investigación Biomédica En Red en Enfermedades Raras (CIBERER); and Fanconi Cancer Foundation.

## Authors’ contributions

M.S., W.C., K.S., J.Su., J.M., A.D. conceived the experiments. MS and WC wrote the main manuscript text. MS and WC prepared the figures. YSG, DChe and JB synthesized the SC4 analogue compounds. CL, JSc and JO sourced the other AMPK activators and provided intellectual input. HB, JL, MS performed high-throughput screening experiments and SR, MS and KC performed analysis. JSu, DC and JM performed micronucleus and G2-arrest assays and had intellectual input into the project. All authors reviewed the manuscript.

## Availability of data and materials

Data generated during this study are available from the corresponding author upon reasonable request and under a data sharing agreement. Access to certain molecular structures contained within the compound libraries screened in this study may require separate approval and/or data sharing agreements with the organisations that deposited those compounds.

## Methods

### Recombinant proteins preparation

Recombinant human biotinylated-avi-ubiquitin, Flag-FANCB, MBP-FANCC, FANCE, FANCF, FANCL, FANCX, FANCT, *X. laevis* GST-FANCD2 and Flag-FANCI were expressed and purified according to what has been previously published^8,12,62^. Human recombinant UBE1 was purchased from Boston Biochem

### FANCX mutant protein preparation

Flag-FANCB, FANCL and FANCX are expressed purified as a subcomplex using the baculovirus system. The baculovirus-ready vector that houses the three genes is cre-recombinated pFL Flag-FANCB eGFP + pSPL FANCL FANCX. Geneblocks containing the mutations were synthesized by IDT and cloned into the vector via a SfiI restriction site in FANCX. Insertion of mutations were confirmed by sanger sequencing. Expression and purification of the mutant FANCB-L-X subcomplex then followed the previously published procedure^12^.

### Compounds for screening

Compounds for the high-throughput screens were sourced from Compounds Australia (Griffith University, Queensland), dissolved in DMSO and screen-ready in a 384-well format. Four sub-libraries were screened from the Compounds Australia collection. These included the Enamine and ChemDiv scaffold library (27,494 compounds), a library of FDA-approved drugs and other academic drug-like molecules (8,067 compounds)^8^, a library from CSIRO (9643 compounds) and an epigenetic and kinase inhibitor library (609 compounds).

A-769662 was purchased from Selleckchem for hit validation. MSG-compounds were synthesized by the team, synthesis procedure is detailed in Ovens et al^35^ and the subsection below. All compounds were dissolved in DMSO at 10 mM, aliquoted and stored at −20°C.

Fatty-acyl CoA compounds were dissolved in deionised water fresh on the day of use at a concentration of 1 mM. Palmitate was dissolved in DMSO due to its low solubility in water.

### Procedures for the preparation of MSG016 and its intermediates

#### Ethyl (1r,4r)-4-((6-chloro-5-iodo-1-((2-(trimethylsilyl)ethoxy)methyl)-1H-imidazo [4,5-b]pyridin-2-yl)oxy)cyclohexane-1-carboxylate (1)

To a solution of ethyl (1*r*,4*r*)-4-hydroxycyclohexane-1-carboxylate (189 mg, 1.10 mmol) in DMF (6.00 mL) was added DBU (380 mg, 2.50 mmol) and 6-chloro-5-iodo-2-(methylsulfonyl)-1-((2-(trimethylsilyl)ethoxy)methyl)-1*H*-imidazo[4,5-*b*]pyridine (488 mg, 1.00 mmol). The reaction was stirred at room temperature for 2 h. The solvent was removed in vacuo, the residue acidified with a 10% aq. citric acid solution and extracted with EtOAc (20.0 mL × 3). The combined organic layers were washed with water, brine, dried (MgSO_4_) and concentrated in vacuo to give the crude material. The product was purified by silica gel chromatography (0-30% EtOAc/ *n*-Hexane) to afford the title compound (325 mg, 56 % yield). ESI-MS: *m/z* = 580.1 [M +H]^+^.

#### Ethyl (1r,4r)-4-((5-(4’-acetamido-2’,3’,4’,5’-tetrahydro-[1,1’-biphenyl]-4-yl)-6-chloro-1-((2-(trimethylsilyl)ethoxy)methyl)-1H-imidazo[4,5-b]pyridin-2-yl)oxy)cyclohexane-1-carboxylate (2)

[1,1’-Bis(diphenylphosphino)ferrocene]dichloropalladium(II) (41.0 mg, 56.0 µmol) was added to a dioxane (9.00 mL)/water(1.00 mL) solution of ethyl (1*r*,4*r*)-4-((6-chloro-5-iodo-1-((2-(trimethylsilyl)ethoxy)methyl)-1*H*-imidazo[4,5-*b*]pyridin-2-yl)oxy)cyclohexane-1-carboxylate **1** (325 mg, 0.560 mmol), *N*-(4’-(4,4,5,5-tetramethyl-1,3,2-dioxaborolan-2-yl)-2,3,4,5-tetrahydro-[1,1’-biphenyl]-4-yl)acetamide (229 mg, 0.672 mmol) and caesium carbonate (274 mg, 0.840 mmol) at rt under a nitrogen atmosphere, the reaction was heated to 80 °C for 3 h. The reaction was cooled to room temperature, concentrated, and then partitioned between water and EtOAc. The aqueous layer was extracted with EtOAc (30.0 mL × 2) and the combined organic phases washed with brine, dried over MgSO_4_ and the solvent removed in vacuo to give the crude material. The product was purified by silica gel chromatography (0-40% EtOAc/*n*-Hexane) to afford the title compound (210 mg, 56 % yield), ESI-MS: *m/z* = 667.3 [M +H]^+^.

#### (1r,4r)-4-((5-(4’-Acetamido-2’,3’,4’,5’-tetrahydro-[1,1’-biphenyl]-4-yl)-6-chloro-1H-imidazo[4,5-b]pyridin-2-yl)oxy)cyclohexane-1-carboxylic acid (MSG016)

TBAF (1.00 M in THF) (5.00 mL, 5.00 mmol) was added to a solution of ethyl (1*r*,4*r*)-4-((5-(4’-acetamido-2’,3’,4’,5’-tetrahydro-[1,1’-biphenyl]-4-yl)-6-chloro-1-((2-(trimethylsilyl)ethoxy)methyl)-1*H*-imidazo[4,5-*b*]pyridin-2-yl)oxy)cyclohexane-1-carboxylate **2** (210 mg, 0.314 mmol) in THF (5.00 mL). The reaction was heated at 80 °C for 3 h. The volatiles were removed in vacuo and the resultant residue dissolved in MeOH (10.0 mL) and a 2.50 M aq. sol. of NaOH (4.00 mL). The solution was stirred at room temperature for 8 h. after which the volatiles were removed in vacuo, the residue taken up in water (10.0 mL) and adjusted to pH 7 with 2.00 M aq. sol. of HCl, then extracted with EtOAc. The combined organic layers were washed with brine, dried over MgSO_4_ and the solvent removed in vacuo. The resultant crude material was purified by silica gel chromatography (0-15% CH_3_OH/CH_2_Cl_2_) to afford the title compound (20.0 mg, 13 % yield, > 95% pure as judged by HPLC) as a light-yellow solid. ^1^H NMR (400 MHz, DMSO) δ 7.97–7.88 (m, 1H), 7.83 (s, 1H), 7.65–7.57 (m, 2H), 7.55–7.45 (m, 2H), 6.24–6.15 (m, 1H), 5.05–4.90 (m, 1H), 3.92–3.80 (m, 1H), 2.52–2.40 (m, 2H), 2.31–2.15 (m, 3H), 2.17–2.05 (m, 1H), 2.02–1.88 (m, 4H),1.82 (s, 3H), 1.69–1.40 (m, 5H). ESI-MS: *m/z* = 509.1 [M +H]^+^.

### Procedures for the preparation of MSG017 and its intermediate

#### Ethyl (1r,4r)-4-((5-(4-(1-acetyl-1,2,3,6-tetrahydropyridin-4-yl)phenyl)-6-chloro-1-((2-(trimethylsilyl)ethoxy)methyl)-1H-imidazo[4,5-b]pyridin-2-yl)oxy)cyclohexane-1-carboxylate (3)

[1,1’-Bis(diphenylphosphino)ferrocene]dichloropalladium(II) (50.0 mg, 0.0679 mmol) was added to a dioxane (9.00 mL)/water (1.00 mL) solution of ethyl (1*r*,4*r*)-4-((6-chloro-5-iodo-1-((2-(trimethylsilyl)ethoxy)methyl)-1*H*-imidazo[4,5-*b*]pyridin-2-yl)oxy)cyclohexane-1-carboxylate **1** (0.300 g, 0.517 mmol), 1-(4-(4-(4,4,5,5-tetramethyl-1,3,2-dioxaborolan-2-yl)phenyl)-3,6-dihydropyridin-1(2*H*)-yl)ethan-1-one (203 mg, 0.620 mmol) and caesium carbonate (252 mg, 0.775 mmol) at rt under a nitrogen atmosphere, the reaction was heated to 80 °C for 3 h. The reaction was cooled to room temperature, concentrated, and then partitioned between water and EtOAc. The aqueous layer was extracted with EtOAc (30.0 mL × 2) and the combined organic phases washed with brine, dried over MgSO_4_ and the solvent removed in vacuo to give the crude material. The product was purified by silica gel chromatography (0-40% EtOAc/*n*-Hexane) to afford the title compound (175 mg, 52% yield). ESI-MS: *m/z* = 653.2 [M +H]^+^.

#### (1r,4r)-4-((5-(4-(1-Acetyl-1,2,3,6-tetrahydropyridin-4-yl)phenyl)-6-chloro-1H-imidazo[4,5-b]pyridin-2-yl)oxy)cyclohexane-1-carboxylic acid (MSG017)

TBAF (1.0 M in THF) (5.00 mL, 5.00 mmol) was added to a solution of ethyl (1*r*,4*r*)-4-((5-(4-(1-acetyl-1,2,3,6-tetrahydropyridin-4-yl)phenyl)-6-chloro-1-((2-(trimethylsilyl)ethoxy)methyl)-1*H*-imidazo[4,5-*b*]pyridin-2-yl)oxy)cyclohexane-1-carboxylate **3** (175 mg, 0.267 mmol) in THF (5.00 mL). The reaction was heated at 80 °C for 3 h. The volatiles were removed in vacuo and the resultant residue dissolved in MeOH (10.0 mL) and a 2.5 M aq. sol. of NaOH (4.00 mL). The solution was stirred at room temperature for 8 h. after which the volatiles were removed in vacuo, the residue taken up in water (10.0 mL) and adjusted to pH 7 with 2 M aq. sol. of HCl, then extracted with EtOAc. The combined organic layers were washed with brine, dried over MgSO_4_ and the solvent removed in vacuo. The resultant crude material was purified by silica gel chromatography (0-20% CH_3_OH/CH_2_Cl_2_) to afford the title compound (20.0 mg, 15 % yield, > 94% pure as judged by HPLC) as a light-yellow solid. ^1^H NMR (400 MHz, DMSO) δ 7.90 (s, 1H), 7.69–7.60 (m, 2H), 7.59–7.50 (m, 2H), 6.32–6.25 (m, 1H), 5.04–4.95 (m, 1H), 4.12–4.09 (m, 2H), 3.75–3.63 (m, 2H), 2.65–2.57 (m, 1H), 2.40-2.21 (m, 3H), 2.09 (s, 3H), 2.05–1.94 (m, 2H), 1.64-1.45 (m, 4H), 1.34-1.25 (m, 1H). ESI-MS: *m/z* = 495.1 [M +H]^+^.

### High throughput ubiquitination assay

The primary and secondary high-throughput screens were performed at the Victorian Centre for Functional Genomics using liquid handling automation. The FANCD2-monoubiquitination assay was performed in 7 µL reactions in 384-well light grey AlphaPlates (Revvity). Compounds were left to bind proteins for 30 mins in a preincubation step before the reaction was started with ATP at a concentration of 1.14 mM. Reactions were left for 90 mins. The anti-GST acceptor and streptavidin donor alphascreen beads were used to detect biotinylated-FANCI conjugated to GST-FANCD2. Further details of the screening assay can be found in Sharp *et al* 2020. In the primary screen 45,784 compounds were analysed in singlets at 20 µM using the FANCD2-monouiquitination assay. Three sets controls were used in each plate which included 24 maximum signal control reactions (DMSO), 16 minimum signal control reactions (no ATP added to reactions) and 24 inhibitor control reactions (UBE1 inhibitor -TAK243). The Z’ was calculated for each plate to demonstrate robustness using the maximum and minimum signal controls. Z-scores were calculated to normalise the data using the maximum signal control. Primary screen hits were determined considering compounds with a z-score less than −2. The secondary screen analysis was performed on 1,332 compounds (∼3%) selected from the primary screen with the FANCD2-monoubiquitination assay in quadruplicates at 20 µM. Secondary screen hits were determined as compounds with average z-score less than −2. 305 compounds were then analysed with the FANCD2-monoubiquitination assay in a 5-point dose response (0.2, 0.62, 2, 6.2 and 20 µM). Area under the curve was calculated from each dose response to rank the hit compounds on potency.

### Western blot of recombinant FANCD2 monoubiquitination assay

FANCD2-monoubiquitination reactions were set up the same as for the high-throughput screening assay except for the fact that the reaction volume was doubled to 14 µL. Reactions were stopped with 4x LDS buffer (Thermo Fisher) and heated at 80°C for 5 mins. Each sample was loaded on to a 4-12% Bolt™ bis-tris polyacrylamide gel and run in MOPS buffer for 40 mins at 165 V. Proteins were then transferred to Immobilon^®^-FL PVDF membrane via wet tank transfer at 100 V for 1 hour. Transfer buffer was 25 mM Tris 192 mM Glycine 10% methanol. After transfer, membrane was blocked in 0.5 % skim milk PBS for 30 mins. Membranes were then probed with Streptavidin DyLight™ 800 conjugated (Rockland Immunochemicals) at a concentration of 1:5000 in 0.5% skim milk PBS for 45 mins to detect biotinylated ubiquitin. Membrane was then washed 4x in PBS and visualised using LiCor Odyssey CLx scanning in the 800 nm channel.

### Human cell culture

HEK293T cells were obtained from Andrew Deans (St. Vincent’s Institute of Medical Research) and RPE *p53*^-/-^ *BRCA1*^-/-^ isogenic pair of cells were obtained from Alan D’Andrea (Dana Faber Cancer Center). HEK293T cells were cultured in high-glucose-supplemented Dulbecco’s modified Eagle’s medium (DMEM) (Sigma-Aldrich, St. Louis), supplemented with 10% (v/v) fetal bovine serum (FBS). RPE cells were cultured in DMEM/Ham’s nutrient mixture F-12 (Sigma) supplemented with 10% FBS. All cell lines were cultured in a humidified incubator at 37°C in the presence of 5% CO_2_.

### *FANCX* KO via CRISPR-Cas9

A *FANCX* knockout cell line was generated in HEK293T cell using lentivirus containing the the vector LentiCRISPRv2 (Addgene #98290) construct with a guide targeting exon 6 (5’ GGCACUGACAAGCCUCAACG). Lentivirus was made in HEK293T cells. Cells were seeded at a density of 4×10^5^ cells in 1 mL suspension in a 6-well plate. After 24 h, a transfection mixture containing 375 ng of psPAX2, 125 ng of pVSV-G, and 500 ng of LentiCRISPRv2-gRNA-FANCX vector was transfected into the cells. The transfection particles were formed with 3 µL of FuGENE® HD in 100 µL of OptiMEM (Gibco, Thermo Fisher, USA) containing the DNA mixture from above. The transfection particles were left to form over 20 mins at room temperature. The transfection mixture was then added into the HEK293T cells. After 48 h and 72 h after transfection, the supernatant was harvested and pooled, then filtered through a 0.45 µm syringe filter, aliquoted and stored at −80°C until further use. The HEK293T cells that were generated with the virus were kept in culture and put under puromycin selection (2 µg/mL) for 48 h. PCR DNA amplification of the suspect CRISPR cut site was performed (oligos 5’ AGTCAAGATGTGGCACCTGG and 5’ CTGTTCTCTAGCACGCAGGT) and sanger sequence. A c.1459_1469insA mutation was found in *FANCX* resulting in a premature stop codon 305 bp downstream (p.N487fsX587). FA pathway activity of FANCX KO was assessed by performing a FANCD2 western blot on cell lysates (Sup. Fig. 4). In brief, 1×10^6^ cells were seeded in T25 flask then treated for 6 hours with 300 nM mitomycin C. Cells were trypsinised and harvested in RIPA buffer with mammalian protease inhibitors. Cells were lysed with 5 seconds of sonication and protein concentration of lysate was estimated by using BCA method. 50 µg of protein were loaded and separated on a 3-8% Tris-acetate NuPage polyacrylamide gel (Invitrogen). Proteins were transferred to Immobilion-FL PVDF membrane (Millipore) using wet-transfer method in 25 mM Tris, 192 mM Glycine and 10% methanol overnight at 30V. Membrane was blocked with 0.5% skim milk in PBS and incubated overnight with rabbit anti-FANCD2 (ab108928, abcam) at a concentration of 1:2000. Membrane was washed in PBS, then incubated for 1 hour with anti-rabbit IgG DyLight800 9 (SA5-35571, ThermoFisher) at a concentration of 1:5000. Membrane was again washed and visualized using LiCor Odessey Clx.

### Cell dose response assay

Hit compounds A-769662 and MSG010 were analysed in a dose response assay to demonstrate synthetic lethality with the *BRCA1* deficiency. Two isogenic cell line pairs were used in the experiment – HEK293T with and without FAAP100 KO and RPE *p53^-/-^*with and without *BRCA1* KO. HEK293T and RPE cells were seeded at 400 cells and 1,500 cells per well of a 384 well plate, respectively. The cells were dosed at 24 h and 72h after seeding with A-769662, MSG010 or Olaparib (synthetic lethal control). The dose range was 0.01 to 10 µM, with 10 concentrations in a 10^1/3^ dilution series. Cells were dosed in triplicate. Growth was observed every 4 h up until the 144h timepoint using IncuCyte SX5 live cell imager (Sartorius).

### FANCD2 foci assay

Nuclear FANCD2 foci were measured in high throughput by quantitating DAPI stained cell nuclei of HCT116 cells using automated confocal imaging (CellInsight CX7 LZR, ThermoFisher Scientific). Control DMSO-treated cells were at 85% confluence after 6 h drug exposure. Drug exposure was performed as a 5-point dose response (0.2-20 µM) in triplicates. FANCD2 foci were detected with immunofluorescence using rabbit anti-human FANCD2 (abcam, ab108928). The secondary antibody used was Zenon™ antirabbit IgG Alexa Fluor™ 647 (ThermoFisher). Foci were detected using Cellomics HCS Studio SpotDetector.V4 BioApplication.

### Micronucleus and G2/M arrest assay

U2OS and FANCA KO cells were used in both the micronucleus and G2/M arrest assays, like previously^28^. Briefly, cells were seeded at 10,000 cells/well of a 96-well plate. Cells were dosed with either 1 µM or 5 µM of A-769662 coupled with diepoxybutane at a concentration of 1 µg/mL, and left for 48 hours. EMA stain was added to cells at a concentration of 0.025 mg/mL and incubated on ice for 30 mins. Cells were washed with PBS + 2% serum and then lysed with 50 µL of lysis buffer 1 (0.584 mg/mL NaCl + 1 mg/mL sodium citrate + 0.3 µL/mL IGEPAL, 1 mg/mL RNase A and DAPI 2 µg/mL). Cells were incubated at 37°C for 1 h in the dark. 50 µl of lysis 2 buffer was added to cells (85.6 mg/mL sucrose, 15 mg/mL citric acid, and 2 µg/mL DAPI) and plate was incubated in the dark for 30 mins. Samples were then analysed by flow cytometry and gated based on size and DAPI signal to be able to count micronuclei. The measure of DAPI staining was also used for the detection of the G2/M arrest, with the 4N DNA-content population serving as an indicator of cell-cycle accumulation at this phase. Experiment was performed five times. Z-scores were calculated using following equation; Z-score = (sample count – mean untreated control count)/standard deviation of untreated control.

---

**Table 1.**
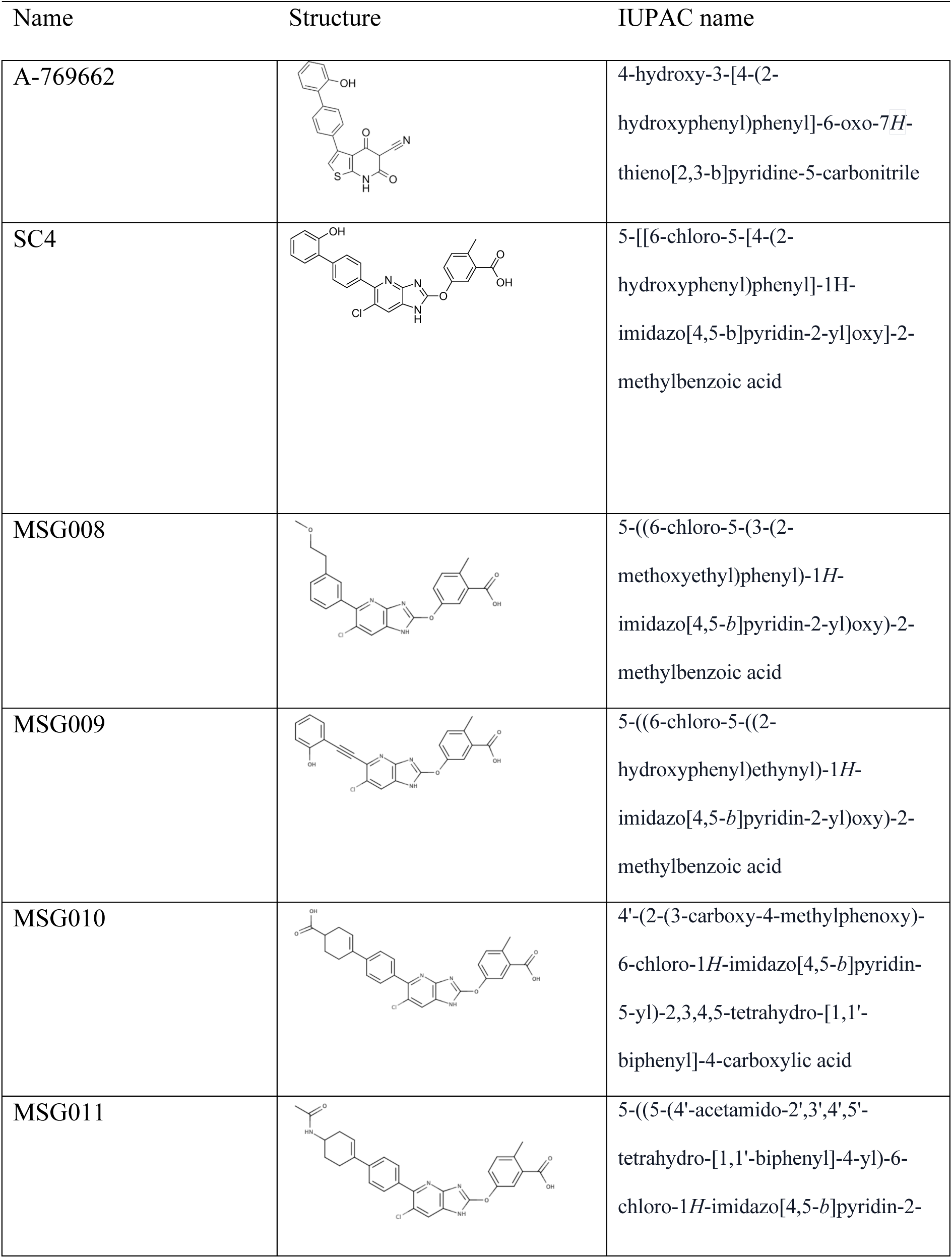
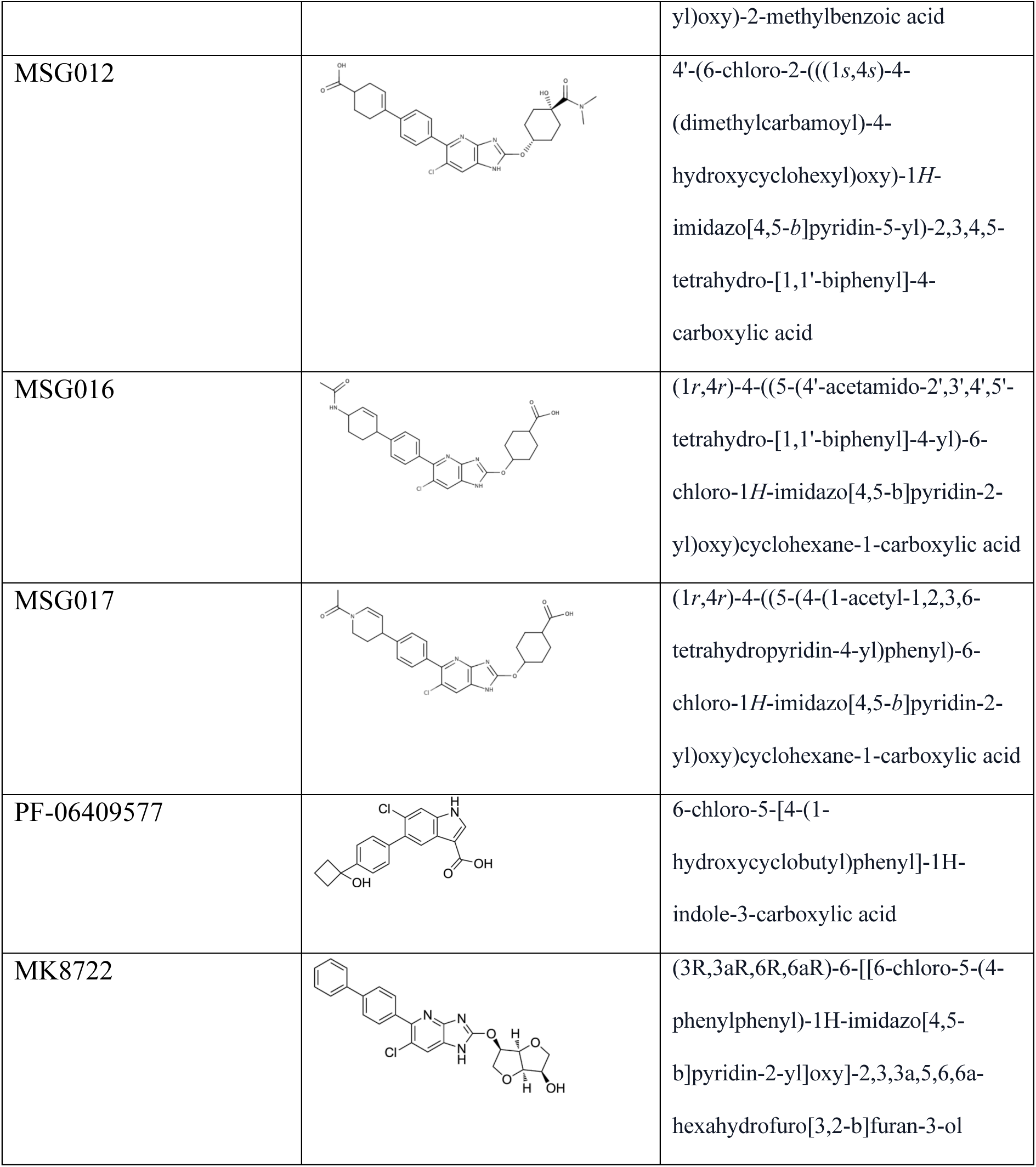
AMPK activating compounds that were screened using FANCD2-monoubiquitination assay.

**Supplementary Table 2.**
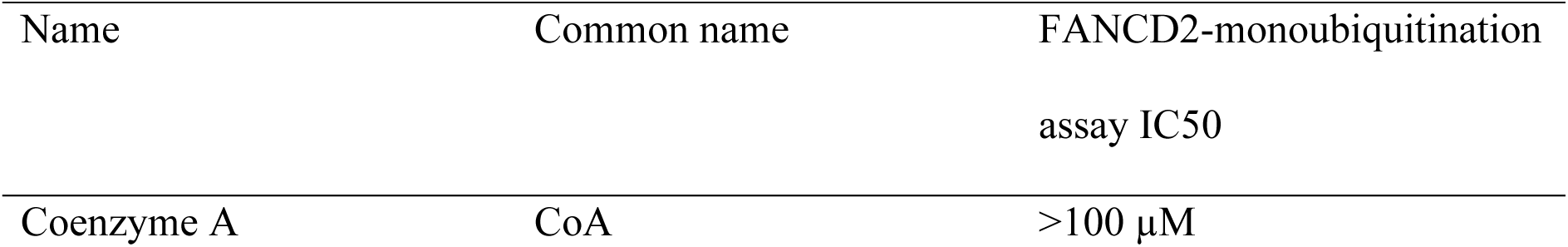
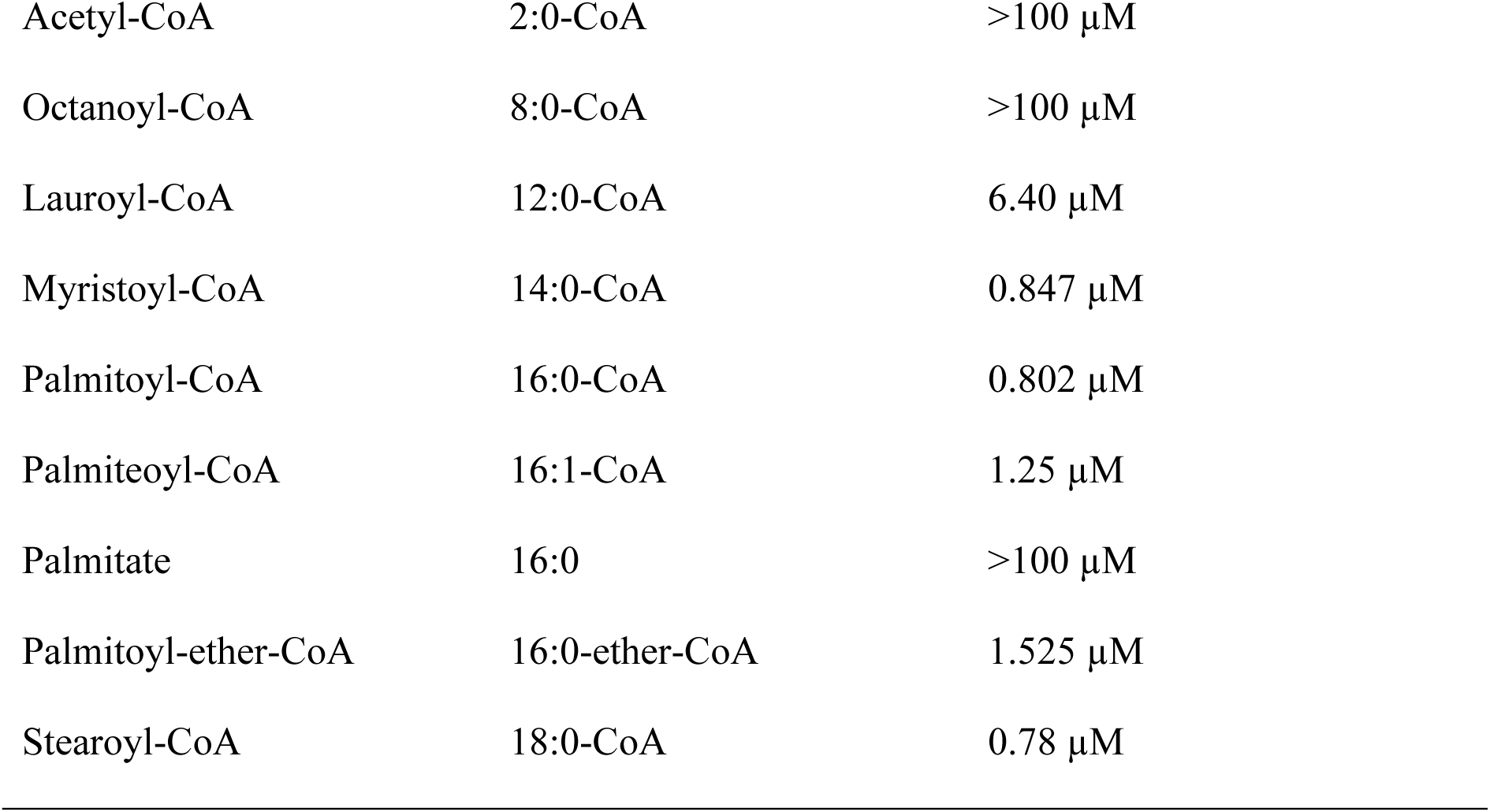
List of fatty acyl-CoA’s, fatty acids and derivatives used in the study.

**Supplementary Figure 1.**
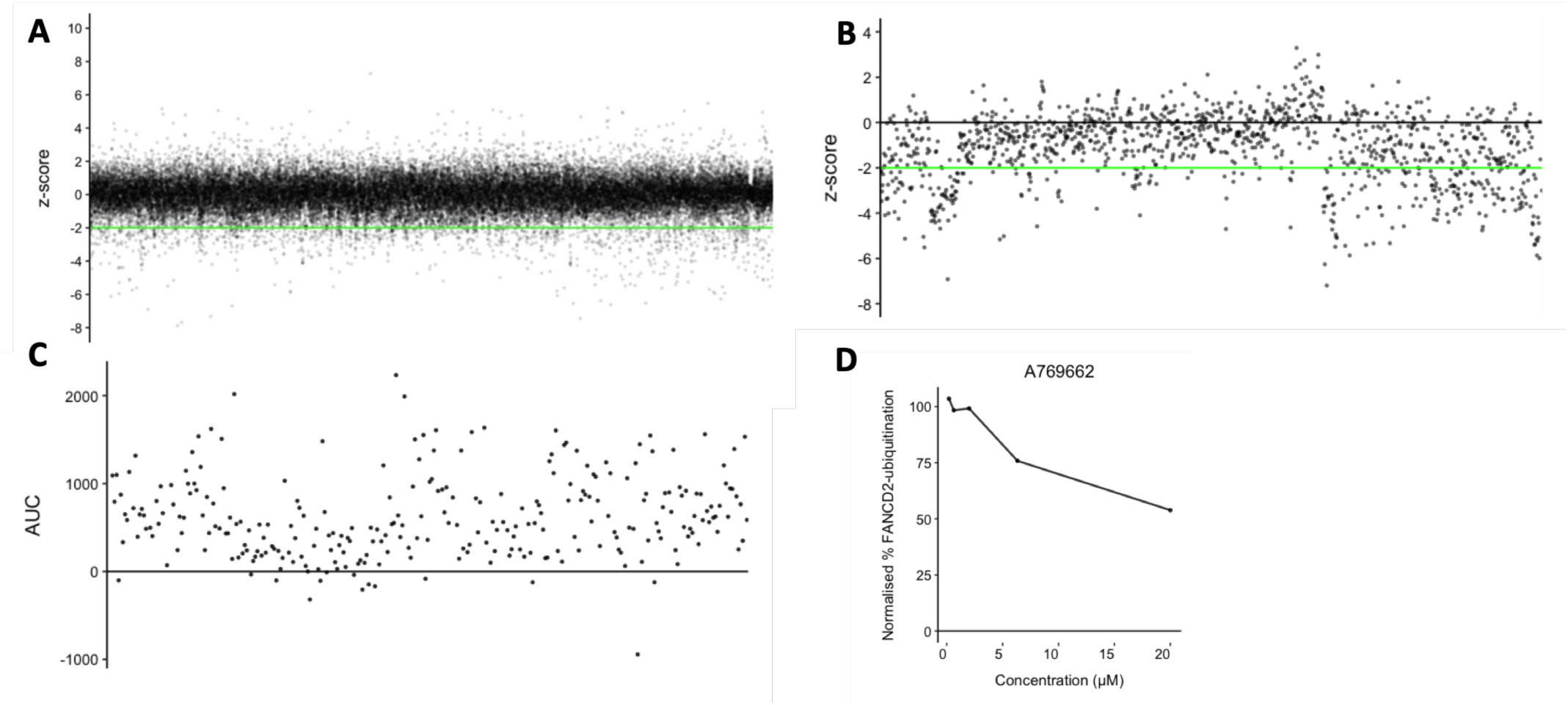
Summary of the high-throughput screen performed with the recombinant FANCD2 monoubiquitination assay. A) Primary screen with 45,784 compounds screened in singlets. Z-score normalisation was calculated for each compound based off the mean signal of the average and standard deviation of the DMSO control. 1,332 compounds had a z-score less than −2 (green line) and were selected for secondary screening. B) Secondary screen of hits displayed as z-scores of the average of quadruplicate data points for each compound in the FANCD2 monoubiquitination assay. 302 compounds had a z-score less than −2 and were selected for dose response screening. C) 302 compounds are ranked on potency from dose response FANCD2 monoubiquitination assay based on the area under the curve (AUC). Normalised percentage inhibition was calculated for each data point of the dose response. High area under the curve represents high potency. D) Hit compound A-769662 inhibitory effect on the percentage of FANCD2 monoubiquitination from the dose response screen.

**Supplementary Figure 2.**
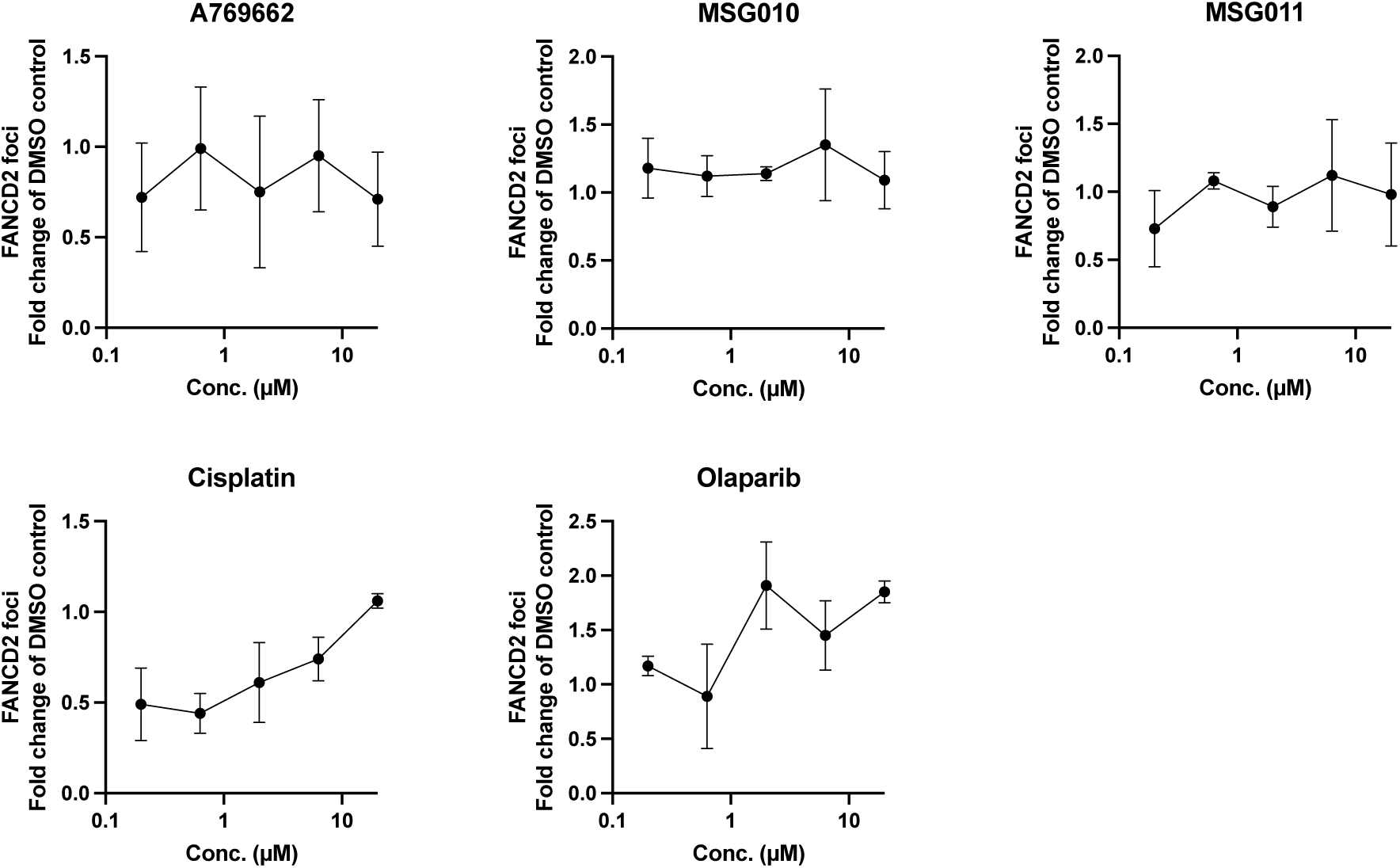
FANCD2 foci assay performed in HCT116 cells. A dose response was performed with three of the AMPK activators as well as controls, cisplatin and olaparib, both of which are expected to increase FANCD2-foci.

**Supplementary Figure 3.**
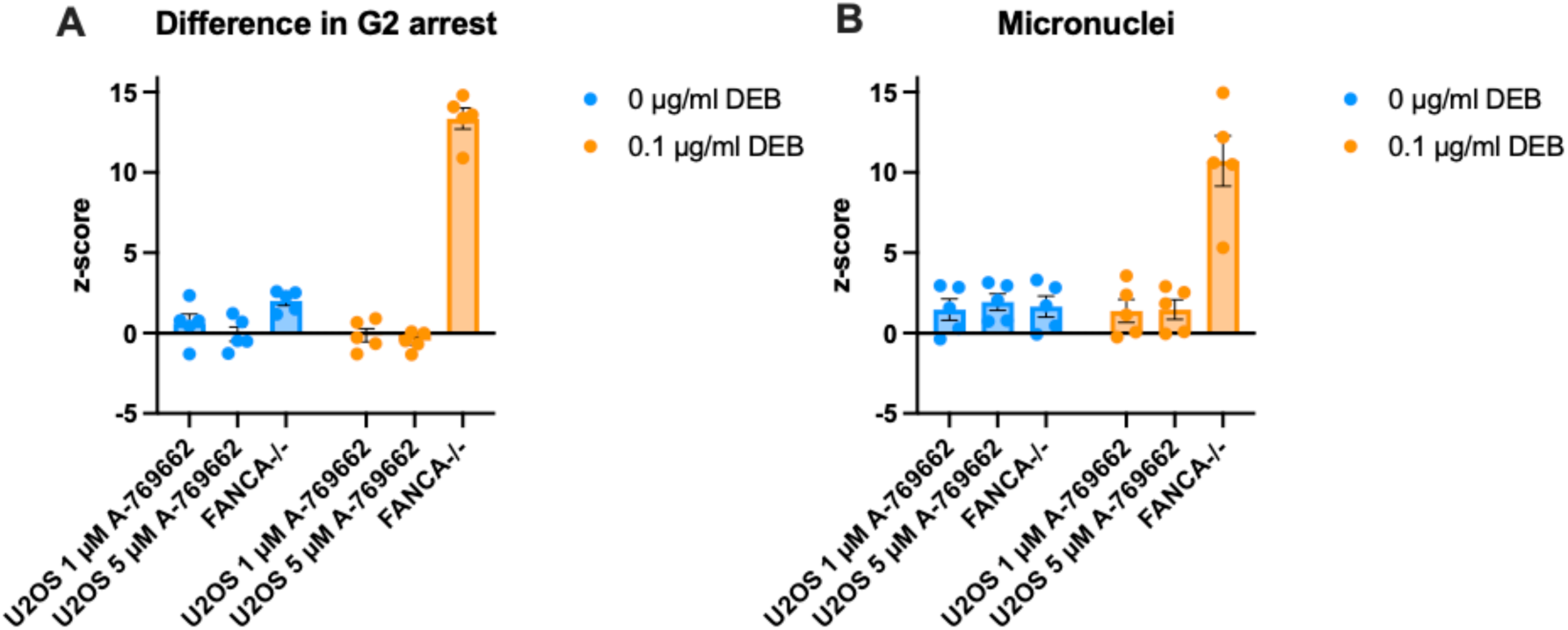
Analysis of FA-like cellular phenotypes with AMPK activator A-769662. FA phenotype control is a *FANCA*^-/-^ cell line. DEB was used to induce DNA damage. A) Analysis of cells which arrested in G2 phase of the cell cycle with A-769662. Data is represented as z-score calculated from the mean and standard deviation of the untreated control (n = 5). B) Micronuclei formation after treatment with A-769662. Data is represented as z-scores calculated from the mean and standard deviation of the untreated control (n = 5).

**Supplementary Figure 4.**
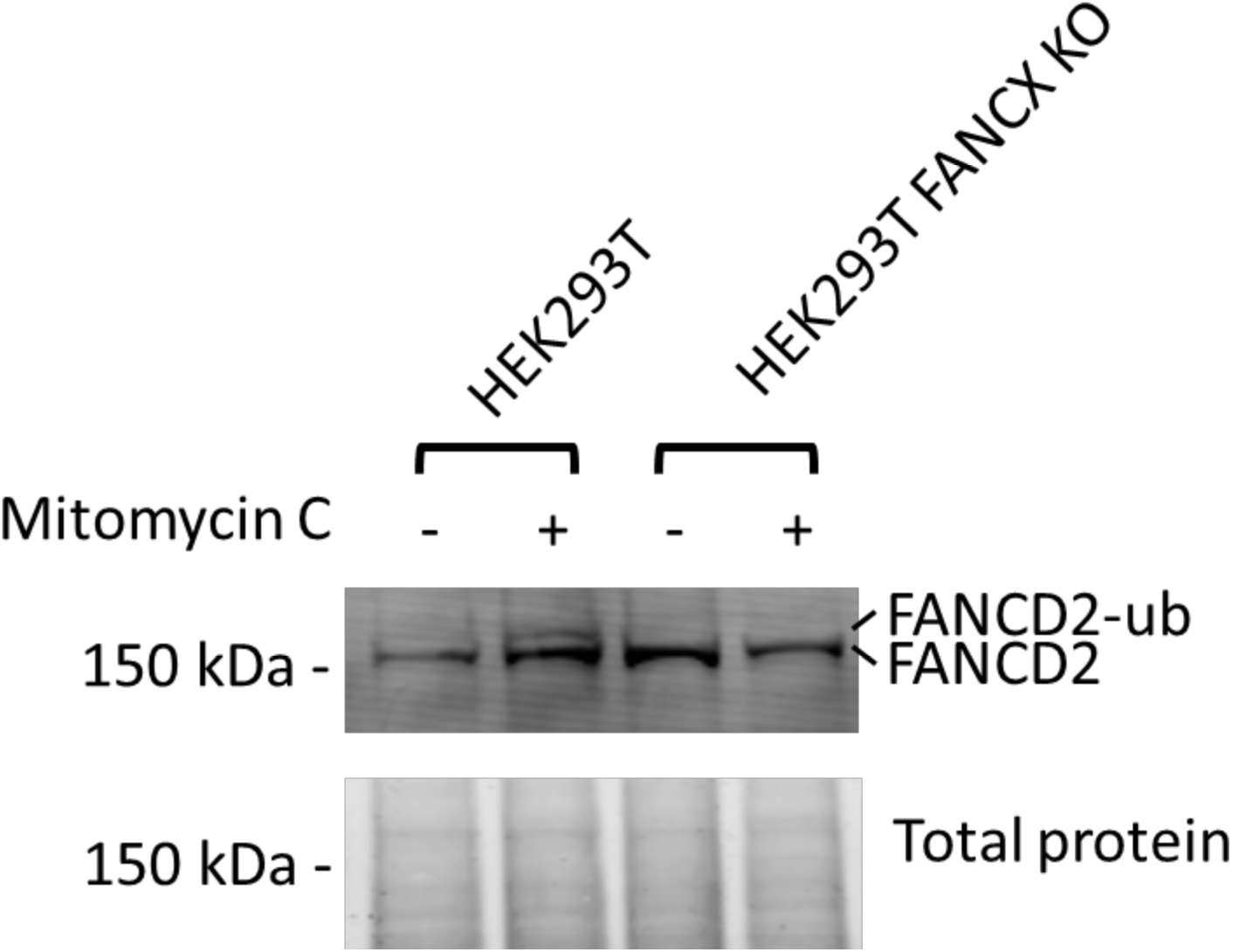
Western blot probing for FANCD2 on HEK293T lysates to demonstrate functional loss of FANCX by observing FA pathway activity. Cells were treated 300 nM mitomycin C to induce FANCD2-monoubiquitination to demonstrate FA pathway activity. Revert™ 700 total protein stain was used as a loading control.

